# Toward Precision Detection of Pyrazinamide Resistance: Critical Concentration Assessment and Rapid Molecular Method Validation

**DOI:** 10.1101/2025.09.03.673939

**Authors:** Yan-Feng Zhao, Li-Li Tian, Nen-Han Wang, Hao Chen, Shuang-Shuang Chen, Lan Yu, Meng-Di Pang, Bei-Chuan Ding, Jie Li, Chuan-You Li, Xiao-Wei Dai

**Author notes:** **Correspondence:** (Chuan-You Li); (Xiao-Wei Dai).

## Abstract

This study aimed to evaluate the diagnostic performance of broth microdilution (BMD) and fluorescence PCR melting curve analysis (MeltPro MTB/PZA assay) for detecting pyrazinamide (PZA) resistance in rifampicin-resistant tuberculosis (RR-TB).RR-TB strains isolated from patients in TB prevention and control institutions and designated hospitals in Beijing from January to December 2009 were analyzed. PZA susceptibility was assessed using BMD, MeltPro MTB/PZ A assay (targeting *pncA* mutations), and whole-genome sequencing (WGS). The sensitivity, specificity, agreement rate, and Kappa value of BMD (at critical concentrations (CCs) of 100 μg/mL and 200 μg/mL)and MeltPro MTB/PZA assay were evaluated using WGS as the reference standard. At CCs of 100 μg/mL and 200 μg/mL, BMD showed resistance rates of 63.6% (70/110) and 45.5% (50/110), respectively. Compared to WGS, BMD demonstrated sensitivities of 96 .1% and 92.2%, specificities of 64.4% and 94.9%, with κ values of 0.590 and 0.872, respectively. The MeltPro assay detected resistance in 44.5% (53/119) of strains, with 89.7% sensitivity, 98.4% specificity, and κ value of 0.882. The MeltPro MTB/PZA assay demonstrates high diagnostic accuracy for PZA resistance screening in RR-TB patients, supporting its utility as a frontline detection t ool. BMD at 200 μg/mL showed better concordance with WGS. Additionally, f ive novel *pncA* mutations (Ile5Thr, Leu27Gln, Ser67stop, Pro69Arg, Trp119Ser) identified in this study expand the catalog of known resistance-associated vari ants and offer new targets for mechanistic investigations.

**IMPORTANCE:** This study evaluates the diagnostic performance of broth microdilution and a rapid molecular assay in detecting pyrazinamide resistance in R R-TB, providing insight into reliable testing approaches and resistance profiles.

## INTRODUCTION

Tuberculosis (TB) remains a leading global infectious cause of death, with an estimated 10.8 million new cases and 1.25 million deaths in 2023. Multidrug-resistant or rifampicin-resistant TB (MDR/RR-TB) accounts for 3.2% of new cases and 16% among previously treated cases worldwide **(1)**. Pyrazinamide (PZA), widely used in both first- and second-line regimens, demonstrates unique sterilizing activity against semi-dormant bacilli. However, PZA resistance has been increasing globally, with particularly sharp rises observed in China—where resistance rates climbed from 8.0% to 11.3% in non-MDR strains between 2 000 and 2010, and from 29.6% to 50% in MDR strains during the same period**(2)**. Understanding PZA susceptibility as early as possible is crucial for guiding the selection and design of effective regimens.

Current phenotypic drug susceptibility testing (pDST) for PZA faces signifi cant limitations. The bactericidal activity of PZA depends on its conversion to pyrazinoic acid (POA) under acidic conditions (pH ∼5.5). However, these acidic conditions, required for PZA activation, concurrently inhibit *Mycobacterium tuberculosis (M. tuberculosis)* growth, rendering the Lowenstein-Jensen proportion method unreliable for PZA susceptibility testing**(3)**. The gold standard for res istance detection is culture-based drug susceptibility testing using BACTEC MG IT 960 system with PZA medium. Nevertheless, this assay has been associated with false-resistant results due to alkalization of the medium caused by a high inoculum size or the presence of bovine serum albumin. Another limitation is t he long turnaround time, as the test requires a primary culture and is often performed on a secondary culture **(4)**. The broth microdilution (BMD) method ca n provide not only qualitative susceptibility results but also the minimum inhibi tory concentration (MIC), which indicats the degree of drug resistance. However, because it requires an acidic environment, PZA cannot be included on the s ame culture plate as other drugs **(5)**. Additionally, neither the World Health Organization (WHO) nor the Clinical and Laboratory Standards Institute (CLSI) h as recommended a PZA-specific critical concentration (CC) for the BMD method, complicating the interpretation of PZA resistance results. These limitations hinder the development and widespread application of the BMD method for detecting PZA resistance.

The 2023 WHO guidelines prioritize whole-genome sequencing (WGS) as the reference method for detecting *pncA* mutations **(6)**, WGS can provide near-complete information by capturing the entire genetic repertoire of a given clini cal *M. tuberculosis* strain **(7)**. However, its cumbersome nature, cost, and techn ical accessibility limit its scalability in high-burden, resource-limited settings **(8)** . Alternative molecular methods, such as targeted *pncA* sequencing (e.g., MeltPro MTB/PZA), demonstrate 94.4% accuracy compared to WGS while being mo re cost-effective and rapid**(9)**.

The present study addresses key challenges in PZA resistance detection by pursuing three objectives: 1) optimizing BMD critical concentrations via WGS calibration for improved resistance quantification; 2) validating the sensitivity a nd specificity of the MeltPro MTB/PZA assay against WGS results; and 3) expanding the catalog of known resistance-associated mutations. The findings aim to support the standardization of PZA susceptibility testing across healthcare set tings.

## MATERIALS AND METHODS

### Subculture of *M. tuberculosis* Clinical Isolates

Clinical isolates of *M. tuberculosis* were retrieved from storage at −80°C a nd thawed at room temperature. A 100 µL aliquot of each bacterial suspension was inoculated onto neutral Löwenstein –Jensen (L-J) medium (Celnovte Biotechnology Co., Zhengzhou, China). The culture tubes were gently rotated to ensure even distribution of the inoculum over the slanted surface. Subsequently, the tubes were incubated at 37°C (Memmert IPP260 incubator, Schwabach, Germany) and observed weekly for colony growth.

### BMD Susceptibility Testing

Pyrazinamide (PZA) susceptibility testing was performed using the *Mycobacterium tuberculosis* Drug Susceptibility MIC Plate (BASO Diagnostics Inc., Zhuhai, China). Several colonies during the logarithmic growth phase (cultures recovered within one month) were selected with a sterile loop. A bacterial suspension equivalent to a 1.0 McFarland standard was prepared in saline using an u ltrasonic homogenizer (BACspreade1100C, TB Healthcare). This suspension was then diluted 1:100 with the kit-provided culture medium. Using an auto-inoculator (BSJ-9612, BASO Diagnostics Inc., Zhuhai, China), 100 µL aliquots of th e diluted inoculum were dispensed into each well of the plate. Both negative control wells (containing 100 µL of drug-free medium) and positive control well s (containing 100 µL of the diluted inoculum without drugs) were included. After inoculation, the plates were sealed and incubated at 37°C. The minimum inhibitory concentration (MIC) was interpreted after 10–14 days of incubation using a dedicated drug susceptibility analysis system (BSP-TB96, BASO Diagnostics Inc.).

### Minimum Inhibitory Concentration (MIC) Determination

The *M. tuberculosis* H37Rv strain (ATCC 27294) was used as the quality control strain for each batch of experiments. A batch was considered valid onl y if the following criteria were met: 1) bacterial growth was observed in the positive control well (drug-free), and 2) no growth was observed in the negative control well (sterility control). If the positive control wells showed no growth or signs of contamination, the experiments were repeated. For the PZA-specific controls, no bacterial growth was observed in the PZA-negative control wells, while growth was present in the PZA-positive control wells (50% and 100%), confirming the validity of the test conditions. The MIC was defined as the lowest drug concentration that inhibited considerable visible bacterial growth com pared to the 50% positive control. The concentration range of PZA in the microplate was 25–800 µg/mL. For result interpretation, critical concentrations (CC) of 100 µg/mL and 200 µg/mL were applied .

### DNA Extraction

Bacterial colonies were harvested during the logarithmic growth phase and transferred to 1.5 mL microcentrifuge tubes containing 1 mL of sterile normal saline. Then, the suspensions were heat-inactivated at 99°C for 10 min and sub sequently centrifuged at 12,000×*g* for 2 min. The resulting supernatant was use d as the template for nucleic acid amplification.

### Meltpro MTB/PZA Assay for Rapid Detection of PZA Resistance in *M. tuberculosis*

DNA was extracted from the colonies using an automated system (LabAid -824S; Zeesan Biotech, Xiamen, China). The MeltPro MTB/PZA assay (MeltPr o® MTB/PZA Test Kit; Zeesan Biotech, Xiamen, China) was then performed according to the manufacturer’s instructions, providing a fully automated process. Briefly, the assay used PCR and melting curve analysis methods in four separ atereaction tubes. It targeted mutations within the 561-bp *pncA* gene and its promoter region (positions -16 to -1) to determine pyrazinamide resistance. A positive assay result confirms the presence of M. tuberculosis complex, and the resistance status of the target gene was classified as resistant, susceptible, or indeterminate. A detailed explanation of the assay results is provided in the supplemental material.

### Whole Genome Sequencing and Bioinformatics Analysis

Extracted genomic DNA was subjected to WGS on an Illumina NovaSeq next-generation sequencing platform. Library preparation was performed using 1 00–500 ng of bacterial DNA with the Illumina DNA Prep Kit (20060059, Illu mina, San Diego, CA, USA) according to the manufacturer’s protocol. Briefly, DNA was fragmented using transposase-mediated tagmentation, followed by the addition of adapter sequences. The resulting fragments were size-selected for an optimal insert length of approximately 300–350 bp, then enriched and quantified. Sequencing was conducted on the Illumina NovaSeq platform using the No vaSeq × Series 1.5B Reagent Kit (300 cycles) (20104705, Illumina, San Diego, CA, USA), generating paired-end reads with an average length of 302 bp. Fo r bioinformatics analysis, raw read quality was assessed using the fastp tool with default parameters. The BWA aligner was used to map the sequences to the Mycobacterium tuberculosis reference genome H37Rv (GenBank accession no. NC_000962.3). Subsequently, the TB-Profiler tool was employed to determine t he spoligotype and to identify mutations associated with resistance to anti-tuberculosis drugs.

### Statistical Analysis

Statistical analysis was performed using SPSS software (version 19.0; IBM, Armonk, NY, USA). Categorical data are presented as rates or proportions. WGS results served as the reference standard for calculating the sensitivity, specificity, agreement rate, and Kappa value of the BMD method and the MeltPro MTB/PZA assay. The strength of agreement based on the Kappa statistic was interpreted as follows: 0.41–0.60, moderate; 0.61–0.80, substantial; and 0.81–1.0 0, almost perfect. Ninety-five percent confidence intervals (95% CIs) were calc ulated using the online tool available at http://vassarstats.net/index.html.

## RESULTS

### Study Selection

The study population included RR-TB isolates from 126 eligible TB patients. These patients were treated in TB prevention and control institutions and designated hospitals in Beijing between January and December 2009. In total, 11 9 isolates were selected for BMD, MeltPro MTB/PZA assay, and WGS to detect PZA resistance. The remaining 5 isolates were excluded because they were subculture negative, had indeterminate MeltPro MTB/PZA assay results, or had insufficient DNA quantity and quality (Figure 1).

**FIGURE 1.**
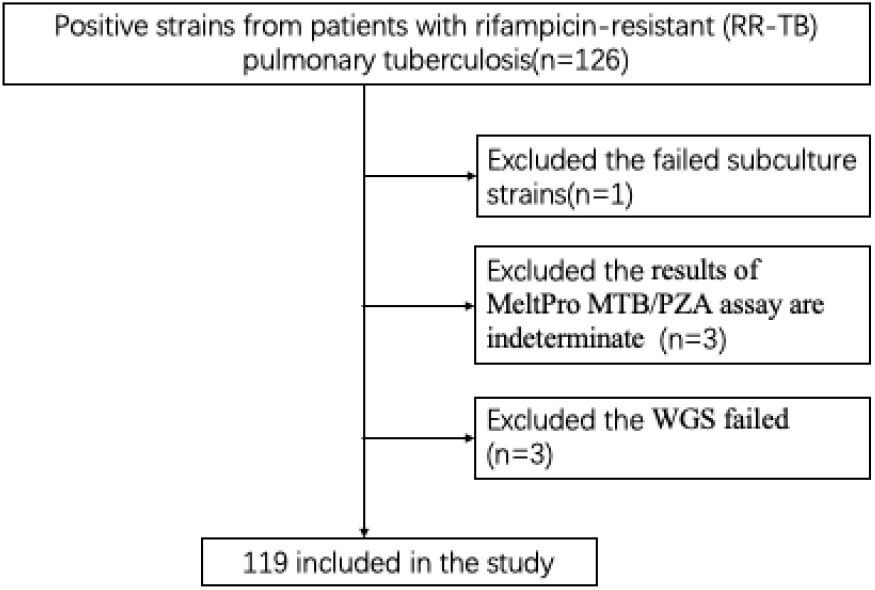
Flow chart of sample collection. Graph show number of isolates/sequences included in the study.

### MIC Distribution and Drug Resistance of PZA

Among the 119 RR-TB strains, 9 strains (7.6%) showed no growth in the PZA positive control wells, and this result was confirmed by repeated tests. There were 11 strains (9.2%) with PZA MICs of 25μg/mL and 50μg/mL respectively, 18 strains (15.1%) with 100μg/mL, and 20 strains (16.8%) with 200μg/m L, 10 strains (8.4%) with 400μg/mL, and 40 strains (33.6%) had a PZA MIC greater than 800μg/mL (Figure 2). After excluding the 9 non-growing positive control strains, among the remaining 110 strains, the PZA resistance rate was 63.6% (70/110) at a CC of 100μg/mL and 45.5% (50/110) at a CC of 200μg/ mL.

**FIGURE 2.**
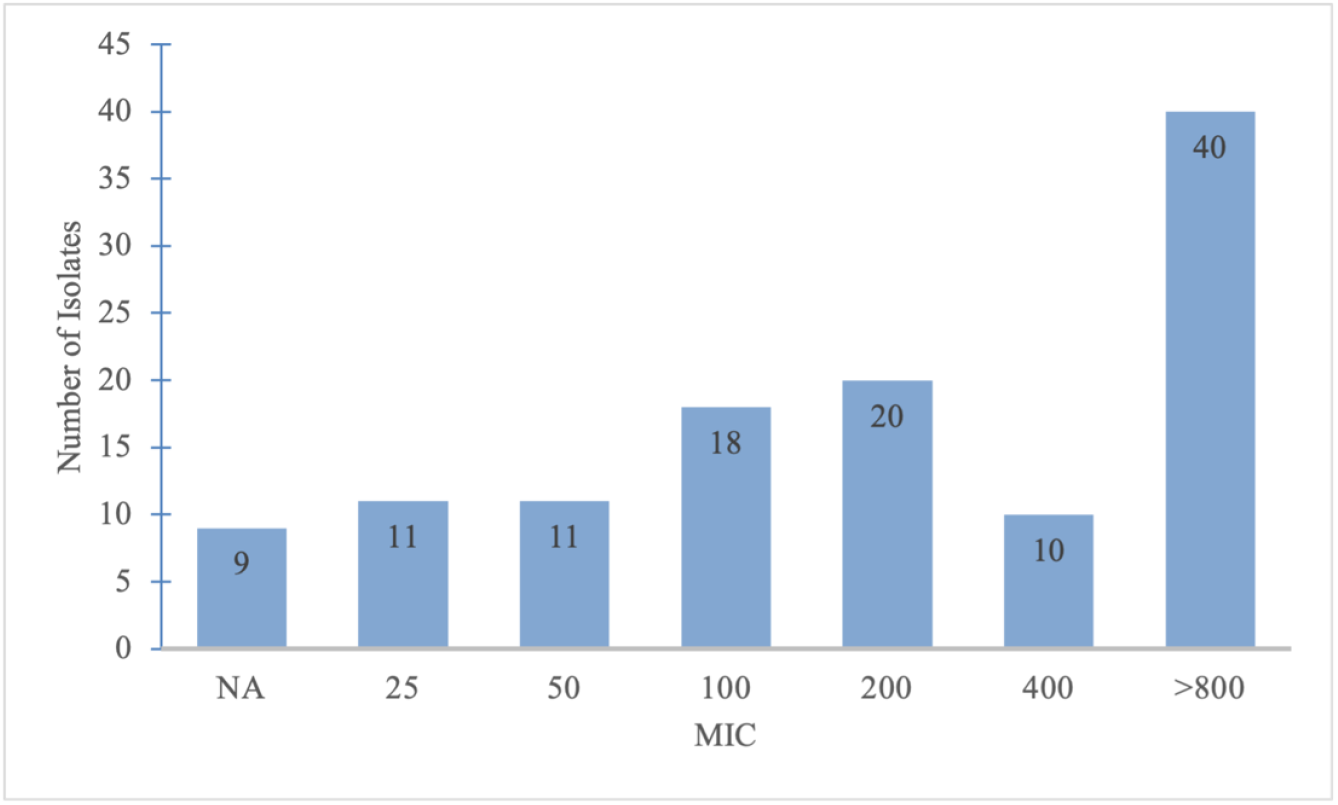
MIC distribution of PZA **Notes: Not Available(NA)**

### Detection of the Meltpro MTB/PZA Assay

Among 119 RR-TB strains, 53 (44.5%) were resistant to PZA. 42 strains had single-channel mutations (18 in Mix A, 13 in Mix B, 11 in Mix C), and 11 strains had multiple-channel mutations (4 in Mix A and Mix B, 4 in Mix A and Mix C, 3 in Mix A, Mix B and Mix C). The mutation frequencies of each channel are provided in Figure 3.

**FIGURE 3.**
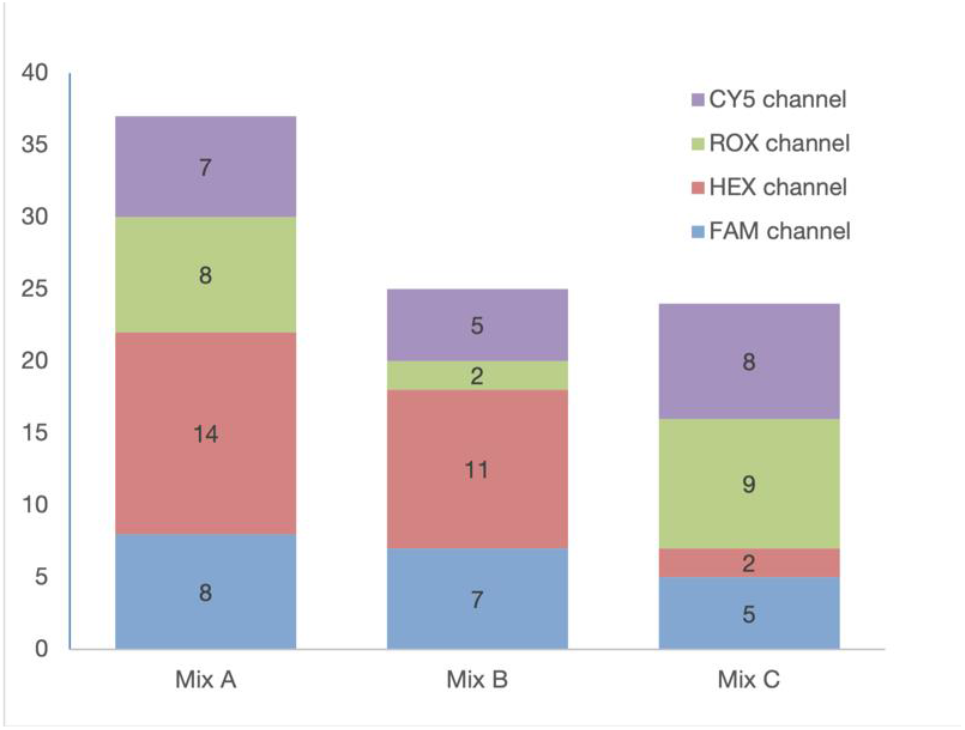
The mutation frequencies of each channel

### Determination of WGS PZA Resistance Mutation

The sequencing results showed that 58 of 119 RR-TB strains had PZA res istance gene mutations, with a mutation rate of 48.7% (58/119). All mutations were found in the *pncA* gene. The mutations were mainly single point mutations, accounting for 60.3% (35/58), while multisite mutagenesis accounted for 6.9 % (4/58). No synonymous mutations were detected. There were 49 mutation types, with the most frequent being A→G at promoter position -11 (4 strains). Among these 4 strains, 3 also carried mutations at other sites. The second and third most common mutations were T→C at codon 281 (Phe94Ser) and A→C at codon 226 (Thr76Pro), each found in 3 strains. Five novel mutations (Ile5T hr, Leu27Gln, Ser67Stop, Pro69Arg, Trp119Ser) were identified. Deletion mutat ions were found in 12 isolates, insertion mutations in 5 isolates, and duplication mutations in 2 isolates. (Table 1).

**TABLE 1.**
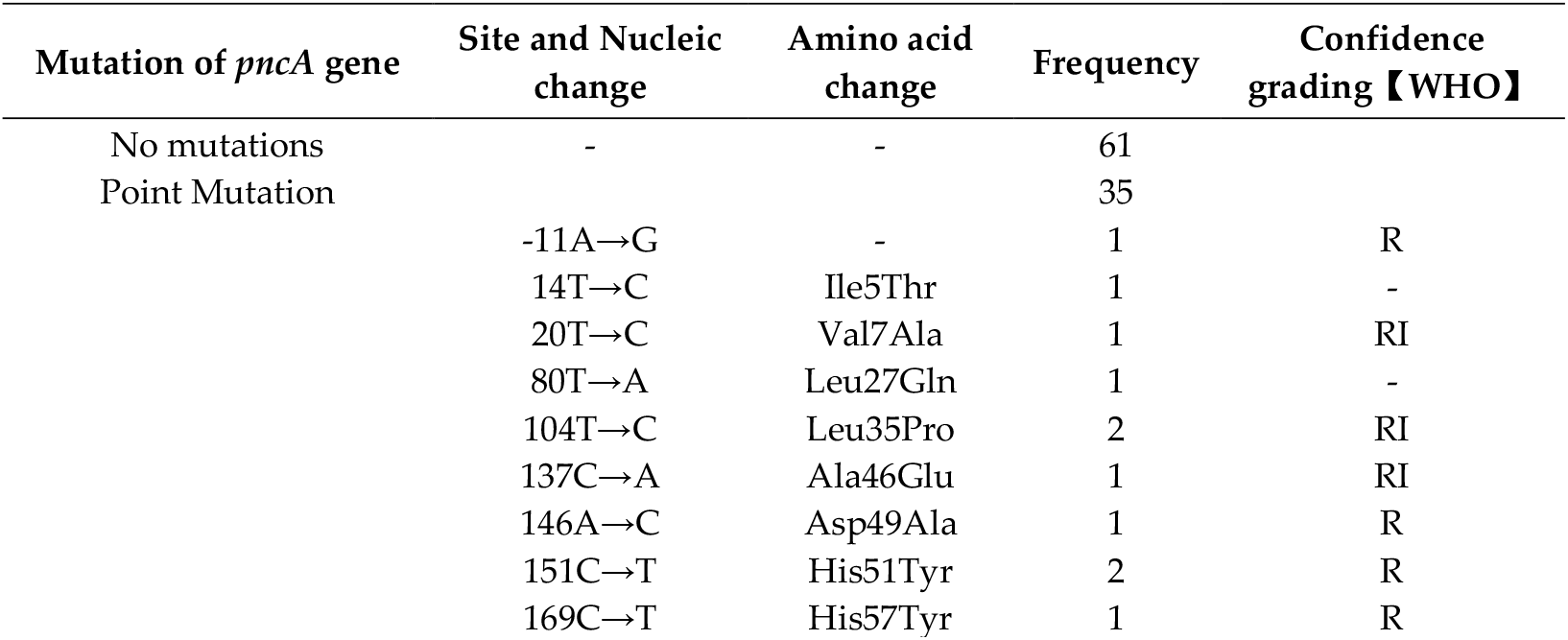
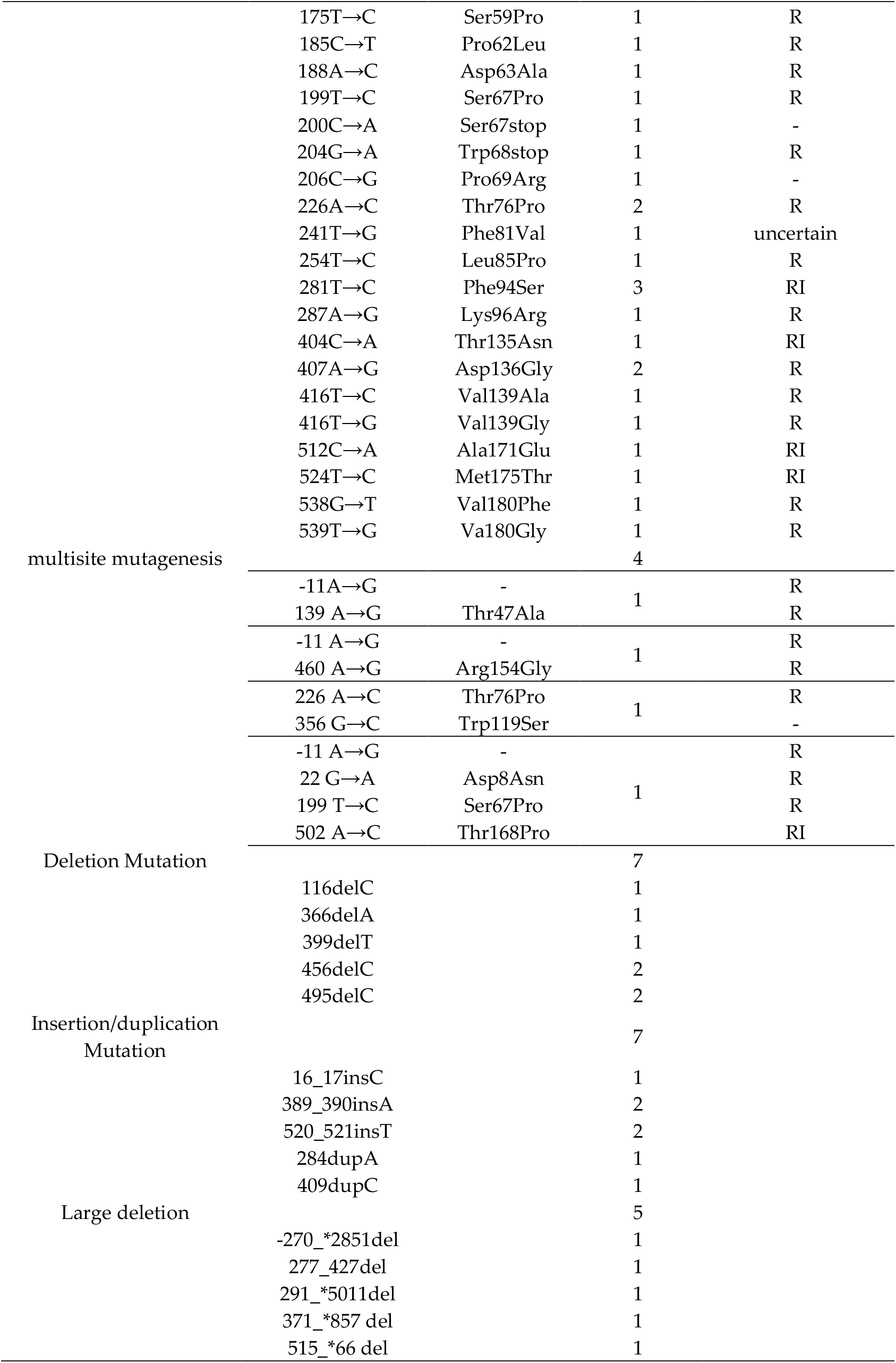
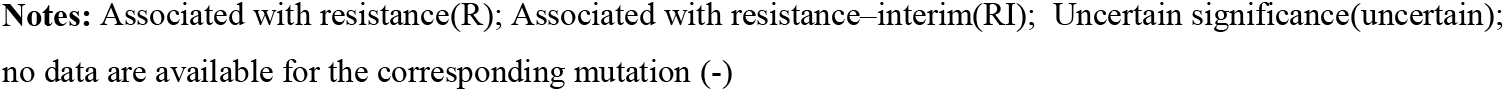
Mutation analysis of *pncA* gene in 119 RR-TB strains

### Performance of the BMD and Meltpro MTB/PZA Assay for Detection of Resistance to PZA

Using WGS as the reference standard, we analyzed 110 cases tested by B MD (CC=100μg/mL and CC=200μg/mL) and 119 cases tested by the MeltPro MTB/PZA assay. The sensitivity of BMD (CC=100μg/mL and CC=200μg/mL) and MeltPro MTB/PZA was 96.1%, 92.2%, and 89.7%, respectively, and the specificity was 64.4%, 94.9%, and 98.4%, respectively. The positive predictive values were 70.0%, 94.0%, and 98.1%, respectively, and the negative predictive values were 95.0%, 93.3%, and 90.9%, respectively. The Kappa value of BMD (CC=100μg/mL and CC=200μg/mL) and MeltPro MTB/PZA were 0.590, 0.872, and 0.882, respectively (Table 2).

**TABLE 2.** Performance of the BMD and MeltPro MTB/PZA assay for detection of resistance to PZA

|  | WGS |  | Sensitivity<br>(%,<br>95%CI) | Specificity<br>(%,<br>95%CI) | PPV<br>(%,<br>95%CI) | NPV<br>(%,<br>95%CI) | Accuracy<br>(%,<br>95%CI) | Kappa<br>Value<br>(%,<br>95%CI) |
| --- | --- | --- | --- | --- | --- | --- | --- | --- |
|  | Wild<br>type | Mu<br>tation |  |  |  |  |  |  |
| BMD(MIC=100µg/mL) |  |  | 96.1<br>(0.8679-0.9892) | 64.4<br>(0.5166-0.754) | 70.0 (0.5846-0.7946) | 95.0<br>(0.835-0.9862) | 79.1<br>(0.7057-0.8564) | 0.590 |
| Sensitive | 38 | 2 |  |  |  |  |  |  |
| Resistant | 21 | 49 |  |  |  |  |  |  |
| Total | 59 | 51 |  |  |  |  |  |  |
| BMD(MIC=200µg/mL) |  |  | 92.2<br>(0.815-0.9691) | 94.9<br>(0.8609-0.9826) | 94.0<br>(0.8378-0.9794) | 93.3<br>(0.8407-0.9738) | 93.6<br>(0.8745-0.9689) | 0.872 |
| Sensitive | 56 | 4 |  |  |  |  |  |  |
| Resistant | 3 | 47 |  |  |  |  |  |  |
| Total | 59 | 51 |  |  |  |  |  |  |
| MeltPro<br>MTB/PZA<br>assay |  |  | 89.7<br>(0.7922-0.9517) | 98.4<br>(0.9128-0.9971) | 98.1<br>(0.9005-0.9967) | 90.9<br>(0.8155-0.9577) | 94.1<br>(0.8836-0.9712) | 0.882 |
| Wild type | 60 | 6 |  |  |  |  |  |  |
| Mutation | 1 | 52 |  |  |  |  |  |  |
| Total | 61 | 58 |  |  |  |  |  |  |

## DISCUSSION

According to WHO estimates, 400,000 people developed MDR/RR-TB globally in 2023, with China reporting 29,000 cases, ranking fourth worldwide **(1)**. PZA is an important component of the standard first-line regimen for TB **(10)** and is also included in WHO consolidated guidelines for treating multidrug-resistant (MDR) and extensively drug-resistant TB (XDR-TB) **(11)**. Recent studies have highlighted the growing challenge of PZA resistance in MDR-TB cases. A 20 23 meta-analysis systematically evaluated this issue by estimating the global weighted pooled resistance (WPR) rate of PZA across WHO regions. Using STATA software for statistical analysis, the authors reported a PZA WPR of 57% (95% CI: 48–65%) in MDR-TB isolate **(12)**. These findings align with regional surveillance data from China(2000–2010) **(2)**. Moreover, after adding PZA to t he treatment regimen, the treatment failure rates were twice as high in patients with PZA-resistant MDR-TB compared to those with PZA-sensitive MDR-TB **(13)**. Timely PZA susceptibility testing for MDR-TB isolates is essential for designing effective regimens and preventing the emergence of extensively drug-resi stant strains. Several studies using the MGIT 960 system reported PZA resistance rates in MDR-TB ranging between 41.01% and 57.7% **(14–19)**. In this study, the BMD method was used to analyze the MIC distribution of PZA resistance among RR-TB isolates. At a CC of 100μg/mL, the PZA resistance rate was 63.6%, while at 200μg/mL, the resistance rate decreased to 45.5%. To determine the optimal critical concentration for PZA resistance detection, we performed WGS on the RR-TB strains in our study. The results demonstrated 93.6% concordance between sequencing data and the 200μg/mL CC, with a Kappa value of 0.872 indicating excellent agreement. These findings indicate that using 200 μg/mL as the BMD critical concentration effectively reduces false-positive results for PZA resistance.

PZA resistance is acquired by *M. tuberculosis* primarily through mutations in the *pncA* gene that diminish the expression of pyrazinamidase /nicotinamidase (PZase) **(20)** or reduce its enzymatic activity **(21,22)**, thereby reducing PZase -mediated conversion of PZA to its active form, POA. In this study, the MeltPro MTB/PZA assay was used to detect mutations associated with PZA resistance. The assay targeted a 561-bp region of the *pncA* gene, including both the resistance-determining and promoter regions (−1 to -16 bp). The results showed t hat 44.5% of RR-TB patients were resistant to PZA, with the MeltPro MTB/P ZA assay demonstrating strong agreement with WGS, yielding a concordance r ate of 94.1% and a Kappa value of 0.882. This consistency was slightly higher than that reported by Li et al. **(23)**, which may be explained by the use of sputum specimens in their study. WGS confirmed that all PZA resistance-conferring mutations were located in the *pncA* gene in this study, likely contributing to the high concordance observed between the two methods. Furthermore, this study analyzed mutation patterns using melting curve channels for PZA detection n. The assay comprised 14 channels distributed across four wells (A, B, C, an d D), with well D serving as the quality control channel. *pncA* gene mutations were identified by comparing sample peaks against 36 wild-type reference peaks across 12 channels in wells A, B, and C of the positive control. Notably, mutations were predominantly localized to the A-HEX, B-HEX, and C-ROX channels. However, as the commercial kit did not specify the nucleotide positions corresponding to individual channels, this study could not delineate the exact base-level mutation sites. In conclusion, the MeltPro MTB/PZA assay represents a rapid, semi-automated molecular platform for the timely detection of PZA resistance in MDR/RR-TB cases.

The WGS results in this study revealed that point mutations in the *pncA* gene accounted for 67.2% of all detected cases, with 49 distinct mutation types identified. The most frequently mutated site was the -11 position in the prom oter region (observed in 4 strains, 6.9%), a prevalence slightly lower than the 7.9% reported in a the nationwide multicenter study by Zhang et al. **(24)**. Basi cally consistent with the results reported by Li et al. **(25)**, who observed a 6.8 % mutation rate at this locus in strains from Zhejiang, Jiangsu, and Sichuan. Mutations in the promoter region were associated with PZA resistance, as demonstrated by Pang Y et al. **(26)**, who showed that such mutations in this region lead to PZA resistance through downregulation of *pncA* gene expression. Overall, the predominant mutation types in the *pncA* gene are largely consistent across different regions of China, though variations in mutation frequencies may occur. These discrepancies could be attributed to factors such as sample size, strain genotype diversity, and local medication practices. In this study, we identi fied frameshift mutations caused by deletions or insertions in both the coding a nd promoter regions of *pncA*. Such mutations are likely to severely impair prot ein function or even result in complete loss of function, thereby contributing to drug resistance. According to the WHO guideline “Catalogue of mutations in *Mycobacterium tuberculosis* complex and their association with drug resistance” (Second Edition) **(27)**, the association between *pncA* gene mutations and drug resistance was clearly established. No mutations categorized as ‘Not associated with resistance’ were identified in this study, One mutation (Phe81Val) classified by WHO as ‘uncertain significance’ was identified in this study. This strain exhibited an MIC value of 400 μg/mL by the BMD method, indicating resistance. Our findings contribute evidence suggesting that this variant may be resistant. Notably, we discovered five previously unreported mutations (Ile5Thr, Leu27 Gln, Ser67stop, Pro69Arg and Trp119Ser),all conferring resistance according to BMD testing .However, their functional mechanisms require further experimental validation.

## CONCLUSIONS

In summary, the BMD test (CC: 200μg/mL) exhibits superior concordance with WGS results for PZA susceptibility testing. The MeltPro MTB/PZA assay targeting the *pncA* gene showed excellent agreement with WGS and could serve as a rapid screening tool for PZA resistance in MDR/RR-TB patients.

This study has several limitations. First, in the BMD assay, growth failure occurred in 9 PZA-positive control wells. This failure might result from decreased bacterial viability due to prolonged storage or excessive subculturing. Another possible explanation is frameshift mutations causing loss of critical protein function. These possibilities require further investigation. Additionally, the Melt Pro MTB/PZA assay can only determine mutation frequencies in specific channels without providing precise mutation site information, precluding detailed com parison with WGS data. Moreover, the drug resistance mechanisms mediated b y five novel mutations (Ile5Thr, Leu27Gln, Ser67stop, Pro69Arg, Trp119Ser) warrant further investigation. Their correlation with phenotypic resistance profiles also needs to be explored.

## SUPPLEMENTARY MATERIALS

Supplemental File 1, PDF File.

## AUTHOR CONTRIBUTIONS

Conceptualization, Yan-Feng Zhao and Xiao-Wei Dai; Methodology, Yan-Feng Zha o, Li-Li Tian, Nen-Han Wang, Hao Chen, Shuang-Shuang Chen, Lan Yu and Xiao -Wei Dai; Software, Meng-Di Pang; Formal analysis, Yan-Feng Zhao and Xiao-Wei Dai; Resources, Jie Li and Xiao-Wei Dai; Writing original draft, Yan-Feng Zhao; Writing review & editing, Yan-Feng Zhao and Xiao-Wei Dai; Visualization, Yan -Feng Zhao; Supervision, Bei-Chuan Ding, Jie Li, Chuan-You Li, Xiao-Wei Dai; Project administration, Chuan-You Li, Xiao-Wei Dai; All authors have read and agreed to the published version of the manuscript.

## INSTITUTIONAL REVIEW BOARD STATEMENT

This study was approved by the Institutional Ethics Review Committee [Appro val Number: 2023 No. (13)].

## INFORMED CONSENT STATEMENT

Not applicable.

## DATA AVAILABILITY STATEMENT

The original contributions presented in this study are included in the article. Further inquiries can be directed to the corresponding authors.

## ABBREVIATIONS

The following abbreviations are used in this manuscript:

BMD: broth microdilution
PZA: pyrazinamide
RR-TB: rifampicin-resistant tuberculosis
WGS: Whole-genome sequencing
CC: critical concentration
TB: Tuberculosis
MDR/RR-TB: multidrug-resistant or rifampicin-resistant TB
pDST: phenotypic drug susceptibility testing
POA: pyrazinoic acid
MIC: minimum inhibitory concentration
WHO: World Health Organization
CLSI: Clinical and Laboratory Standards Institute
CI: confidence interval
XDR: extensively drug-resistant TB
PZase: pyrazinamidase / nicotinamidase

